# Mathematical formulation and application of kernel tensor decomposition based unsupervised feature extraction

**DOI:** 10.1101/2020.10.09.333195

**Authors:** Y-h. Taguchi, Turki Turki

## Abstract

In this work, we extended the recently developed tensor decomposition (TD) based unsupervised feature extraction (FE) to a kernel-based method through a mathematical formulation. Subsequently, the kernel TD (KTD) based unsupervised FE was applied to two synthetic examples as well as real data sets, and the findings were compared with those obtained previously using the TD-based unsupervised FE approaches. The KTD-based unsupervised FE outperformed or performed comparably with the TD-based unsupervised FE in *large p small n* situations, which are situations involving a limited number of samples with many variables (observations). Nevertheless, the KTD-based unsupervised FE outperformed the TD-based unsupervised FE in non *large p small n* situations. In general, although the use of the kernel trick can help the TD-based unsupervised FE gain more variations, a wider range of problems may also be encountered. Considering the outperformance or comparable performance of the KTD-based unsupervised FE compared to the TD-based unsupervised FE when applied to *large p small n* problems, it is expected that the KTD-based unsupervised FE can be applied in the genomic science domain, which involves many *large p small n* problems, and, in which, the TD-based unsupervised FE approach has been effectively applied.

## Introduction

Recently, the tensor decomposition (TD) based and principal component analysis (PCA) based unsupervised feature extraction (FE) (Taguchi, 2020) approach was developed to identify a limited number of genes in *large p small n* problems involving a small number of samples (*n*) with a large number of features (genes) (*p*). This approach, applied to genomic science applications, which frequently involve *large p small n* problems, could successfully identify the limited number of biologically reliable genes that cannot be selected using conventional statistical test based feature selection methods. Nevertheless, because the TD and PCA based unsupervised FE do not include any tunable parameters owing to their linearity, the methods cannot be modified or optimized in failure scenarios. Thus, it is desirable to extend the TD and PCA based unsupervised FE to include nonlinearity.

To this end, we aimed at extending the TD-based unsupervised FE to incorporate the kernel trick (Schölkopf, 2000) and introduce nonlinearity. Because tensors do not have inner products that can be replaced with nonlinear kernels, we incorporate the self-inner products of tensors. In particular, the inner product is replaced with nonlinear kernels, and TD is applied to the generated tensor including nonlinear kernels. In this framework, the TD can be easily “kernelized.” Several researchers have attempted to apply kernel methods to process tensor data. For instance, Signoretto et al. (2011); Signoretto et al. (2012); Zhao et al. (2013a,b) input tensors into vectors (or matrices), which were then used to construct the kernels. However, such conversion may destroy the structural information of the tensor data. Moreover, the dimensionality of the resulting vector is typically extremely high, which leads to the curse of dimensionality and small sample size problems (Liu et al., 2015; Yan et al., 2007). He et al. (2017) proposed an implementation in which the tensor was reproduced using a kernelized TD.

It is expected that these problems can be avoided by computing the inner product of the tensors, as realized in the proposed approach.

## Results

### Mathematical formulations

Suppose there exists a tensor *x*_*ijk*_ ∈ ℝ^*N*×*M*×*K*^, which represents the value of the *i*th feature of the samples with properties *j* and *k*. For example, *x*_*ijk*_ might represent the price of product *i* that a person with a gender *k* and age *j* previously bought. In this case, *N* is the number of products available, *M* is the age, and *K* ∈ [1, 2] represents male or female. Alternatively, in genomic science applications, which are the focus domain of this work, *x*_*ijk*_ is the expression of the *i*th genes of the *k*th tissue of the *j*th person; in this case, *N* is the number of genes, *M* is the number of participants in the study, and *K* is the number of tissues for which the expression of genes is to be measured. As another example, *x*_*ijk*_ might represent the electric current of the *i*th circuit at the *j*th temperature and *k*th atomic pressure; in this case, *N* is the number of circuits in a machine, and *M* and *K* denote the number of recordings of the temperature and atomic pressure, respectively.

The aim of TD-based unsupervised FE is to select the limited number of critical features among all the features (as many as *N*). In the first case, when *i* represents a product, the purpose of analysis may be to identify the limited number of products, the buyers of which are restricted to specific ages and genders. In the second case, when *i* represents a gene, the purpose of analysis may be to determine the limited number of genes whose altered expression may cause diseases. In the third example, when *i* is a circuit, the purpose of analysis may be to identify the limited number of circuits whose malfunctioning may result in the failure of the machine. Thus, in general, the purpose of TD-based unsupervised FE is to screen a small number of significant features from the larger set of features.

In this regard, in TD-based unsupervised FE, higher order singular value decomposition (Taguchi, 2020) (HOSVD) is applied to *x*_*ijk*_ to yield

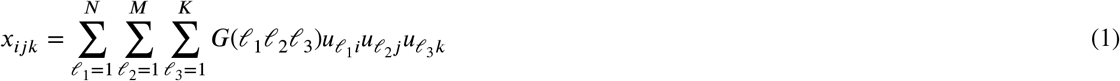

where *G*(*𝓁*_1_*𝓁*_2_*𝓁*_3_) ∈ ℝ ^*N*×*M*×*K*^ is a core tensor, which represents the weight of product 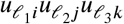 of the contribution to 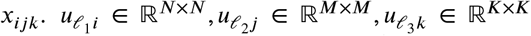 denote singular-value vectors that represent the various dependencies of *x*_*ijk*_ on *i, j, k*, respectively; these vectors are orthogonal matrices.

To select the critical *i*s, we must specify 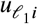 to select *i*s; The *i*s must be those for which the absolute values of 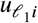 are large. In this context, we must first identify 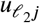 and 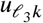, which represent the properties of interest. In the first example, he absolute value of 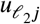, which corresponds to the age, must be larger than a specific age.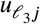, which corresponds to the genders, must be assigned distinct values for the two genders. In the second example,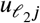, which corrresponds to study participants, must be assigned distinct values for the patients and healthy controls.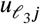, which corrresponds to tissues, must have distinct values for the different tissues. In the third example, 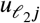 and 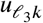 must be assigned extremely large values at specific temperatures and atomic pressures.

After identifying the 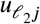 and 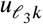 of interest, we attempt to identify *G*(*𝓁*_1_ *𝓁*_2_ *𝓁*_3_) that have larger absolute values en *𝓁*_2_ and *𝓁*_3_. Once *𝓁*_1_ with *G* having the large absolute values is specified, we attempt to identify *i* with large absolute values of 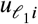. These *i*s might represent the critical products bought by persons with specific ages and gender, genes expressive in certain specific tissues of patients, and circuits in which the electric current increases drastically at specific temperatures and atomic pressures.

The application of TD-based unsupervised FE to problems in genomic science yielded satisfactory results even when conventional feature selection methods based on statistical tests failed (Taguchi, 2020; Taguchi and Turki, 2020; Ng and Taguchi, 2020). Nevertheless, TD-based unsupervised FE has certain limitations. Specifically, owing to the tunable parameters, this approach cannot be modified or optimized in failure scenarios. The proposed method aims at extending the TD-based unsupervised FE to be incorporated with kernel tricks (Schölkopf, 2000). In this scenario, because many kernels can be selected, a better strategy applicable to the target problems can be identified.

To apply the kernel trick, TD must be suitably modified. First, we consider the partial sum as

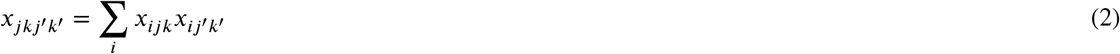

*x*_*ij*′*k*′_ is equivalent to *x*_*ijk*_ with replacing *j, k* with *j*^′^, *k*^′^. Substituting Eq. (1) into the aforementioned equation yields

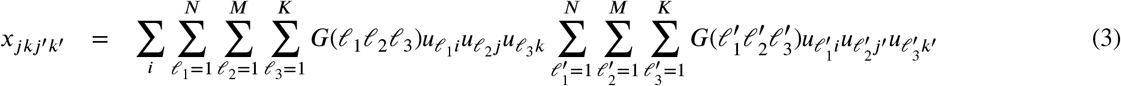

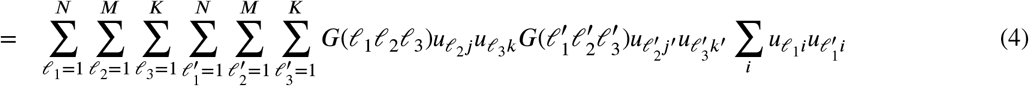

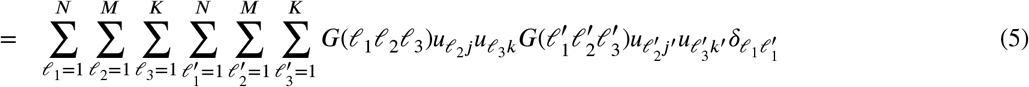

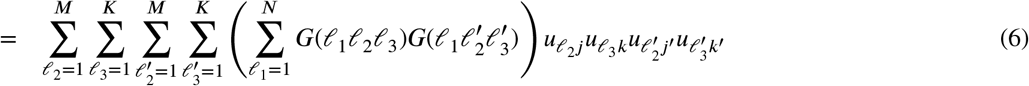

Thus, we can obtain 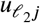and 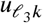 by applying the HOSVD to Eq. (2) as

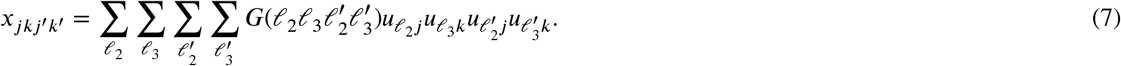

Note that when using the linear kernel, as indicated in Eq. (2), the KTD is equivalent to the (linear) TD.

Because Eq. (2) is expressed in the form of an inner product, it can be easily extended to (nonlinear) kernels. Although in the presented analysis we only considered the radial base function (RBF) kernel as an example, any other kernel can be used in place of the RBF kernel. Eq. (2) can be easily extended to the RBF kernel as

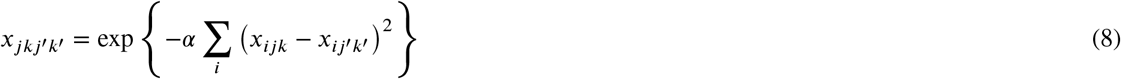

to which the HOSVD can be applied as is. This operation generates an expression corresponding to Eq. (7) with distinct *G*s and 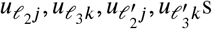.

To integrate two matrices *x*_*ij*_ and *x*_*kj*_ that share the sample *j*, we consider

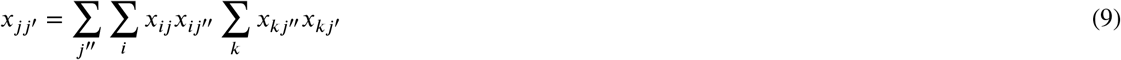

to which the singular value decomposition (SVD) can be applied as is

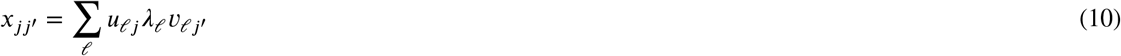

This expression can be easily extended to the RBF kernel as

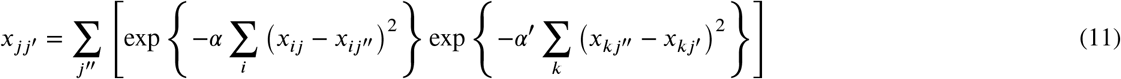

to which the SVD can be applied as is, and an expression similar to Eq. (10) can be obtained with distinct *u*_*𝓁j*,_ *λ*_*𝓁*_, *v*_*𝓁j*_′ Finally, if two matrices, *x*_*ij*_ and *x*_*ik*_, share feature *i*, we have

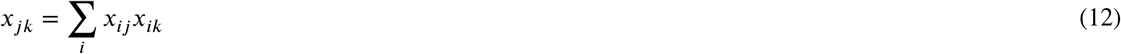

to which the SVD can be applied as is, yielding

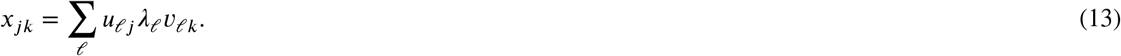

This expression can be easily extended to the RBF kernel as

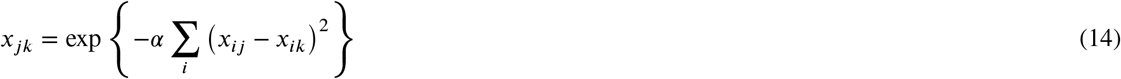

to which the SVD can be applied as is, and we can obtain an expression similar to Eq. (13) with distinct *u*_*𝓁j*,_ *λ*_*𝓁*_, *v*_*𝓁k*_s.

### Application to various data sets

The KTD-based unsupervised FE was applied to various datasets (Fig. 1).

**Figure 1:**
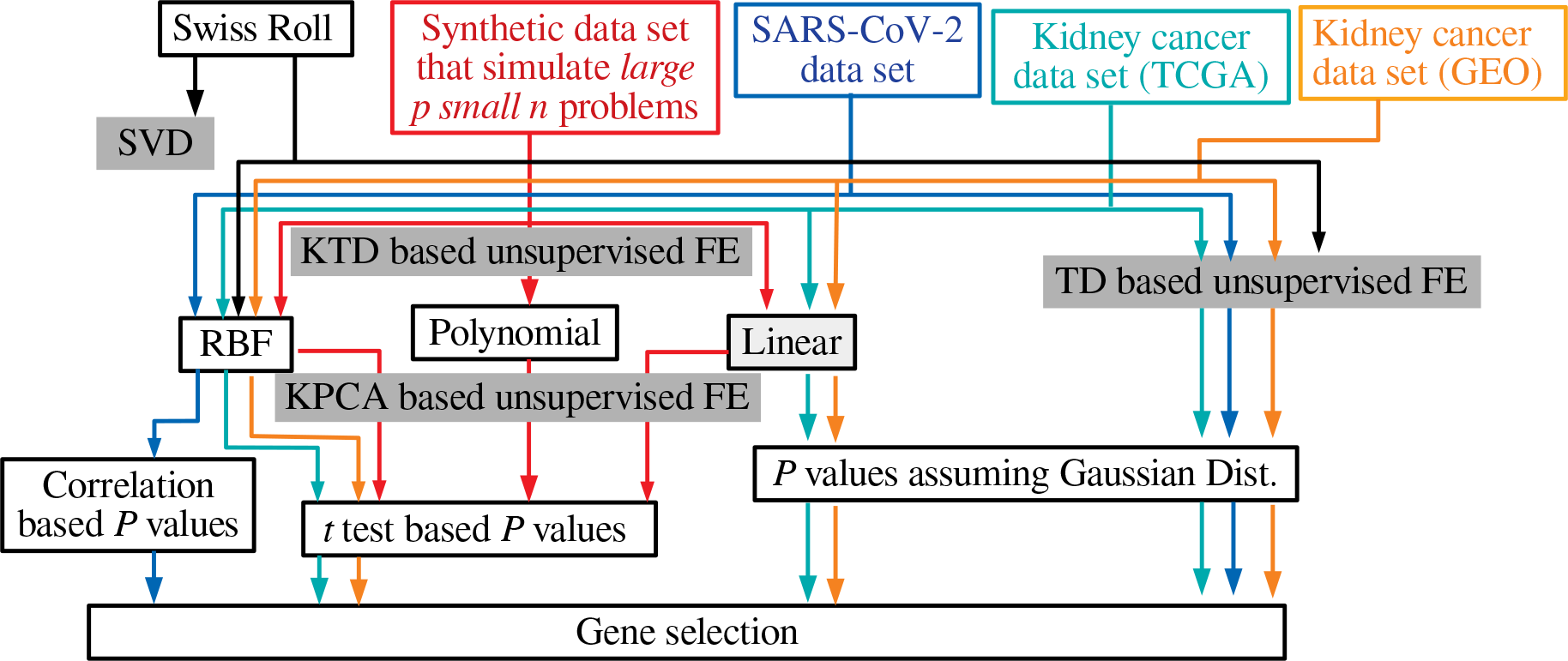
Overview of performed analyses

#### Swiss Roll

To verify if the TD extended to the kernel (KTD approach) can capture the nonlinearity, the Swiss Roll framework is considered, represented as *x*_*ijk*_ ∈ ℝ^*N*×3×10^

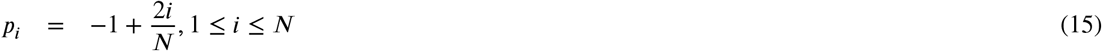

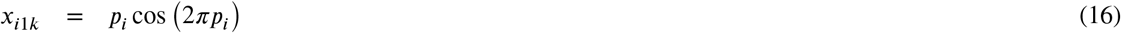

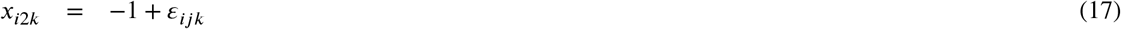

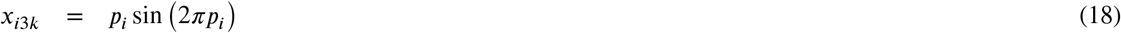

where *ε*_*ijk*_ is drawn the from uniform distribution between −1 and 1 (*x*_*ij*1_ is shown in Fig. 2(A)). Here, Σ_*j*_ *x*_*ijk*_ = 0, Σ_*i*_ *x*_*ijk*_ =*N, N* = 1000. This expression is equivalent to that of 10 ensembles of Swiss Rolls, each of which are generated with distinct *ε*_*ijk*_. To verify that the linear method cannot capture nonlinear structures (i.e., the order of *i* along the curved space) of the Swiss Roll, we apply the SVD to the *x*_*ij*1_ shown in Fig. 2(A). Figure 2(B) shows *u*_*𝓁i*_, 1 ≤ *𝓁* ≤ 3. Clearly, the SVD cannot capture the nonlinear structure of the Swiss Roll. Moreover, even when the HOSVD is applied to *x*_*ijk*_, the nonlinear structure is not well captured (Fig. 2(C)). The kernel based tensor can be generated as

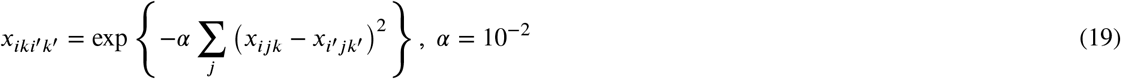

to which the HOSVD can be applied to yield

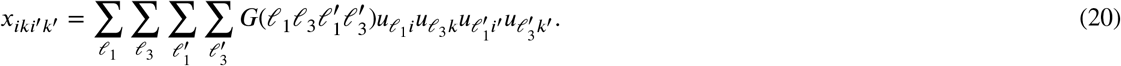

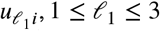 represents the nonlinear structure to a certain extent (Fig. 2(D)). Thus, by replacing the inner product of the tensor with kernels, the developed TD can identify the nonlinear structure of the Swiss Roll, at least partially.

Although in the aforementioned synthetic example, the KTD-based unsupervised FE can successfully identify the nonlinear structure of Swiss Roll, which cannot be captured by the original (linear) TD-based unsupervised FE, the key intent of the TD-based unsupervised FE is to realize the feature selection in *large p small n* problems, with *p ≫ n*. Thus, we must examine if the KTD-based unsupervised FE can outperform the TD-based unsupervised FE in *large p small n* problems.

**Figure 2:**
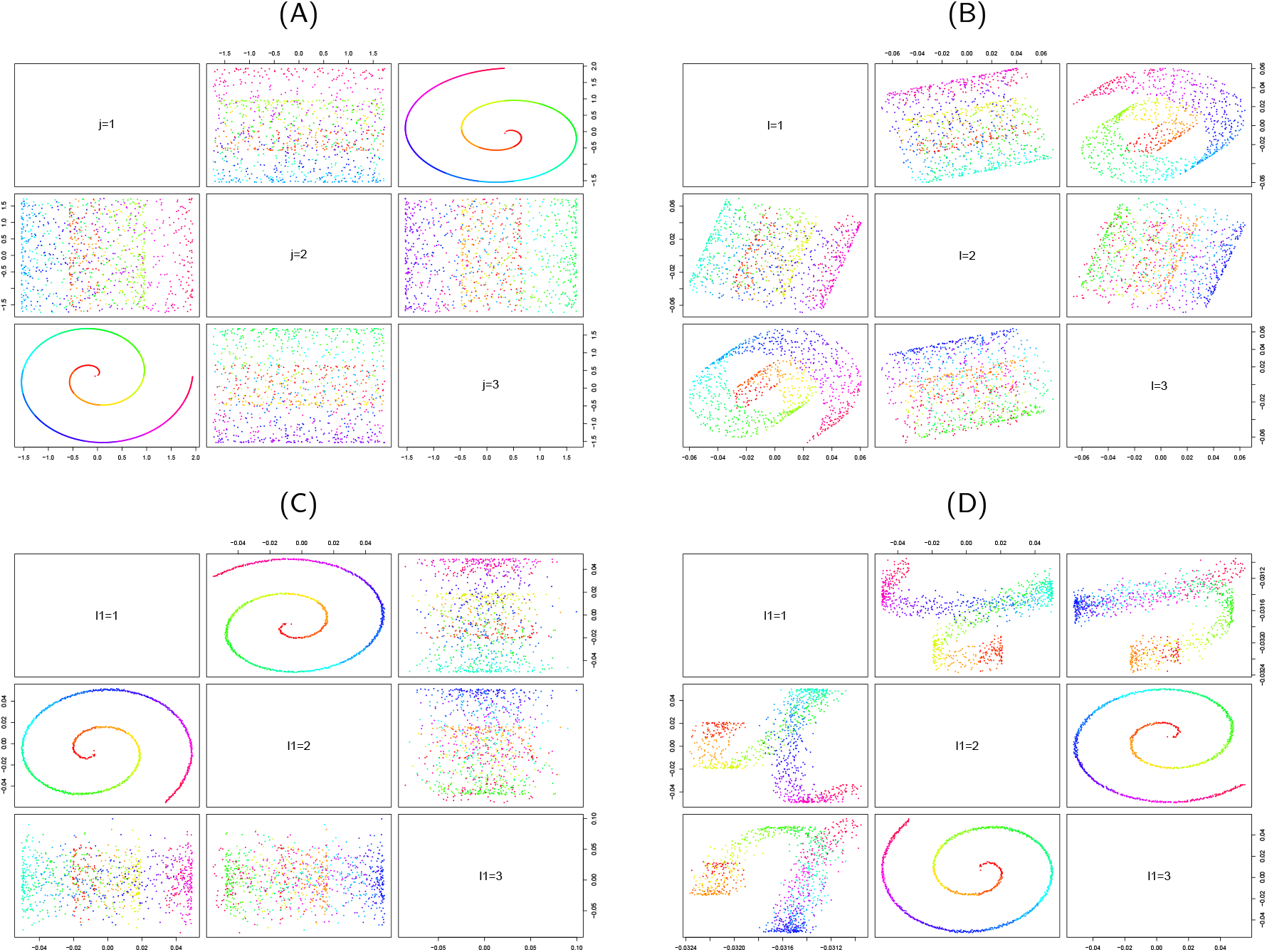
Swiss roll. The colors represent the direction of *i* (1 to N) in gradation. (A) *x*_*ij*1_, (B) *u*_*𝓁i*_, 1 ≤ *𝓁* ≤ 3 by SVD, (C) 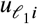, 1 < *𝓁* < 3 by HOSVD, (D) 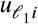, 1 < *𝓁* < 3 by kernel (RBF) based HOSVD.

#### Large p small n problem

We consider the following synthetic example: *x*_*ijk*_ ∈ ℝ^*N*×*M*×*M*^, as

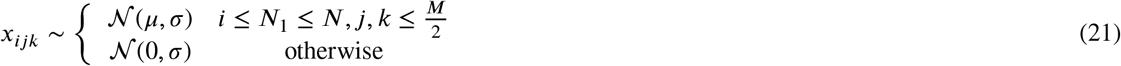

where. 𝒩 (*µ, σ*) is the Gaussian distribution of the mean *µ* and standard deviation *σ*. This problem is slightly challenging as it is a two-class problem although it appears to be a four-class problem. We apply the KTD as well as the kernel PCA (KPCA) as

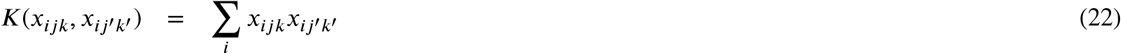

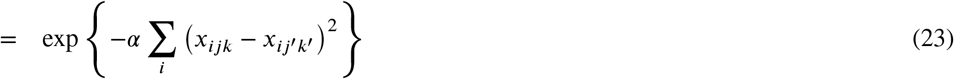

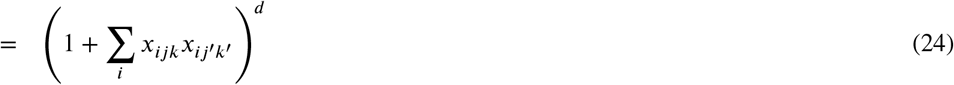

The expressions correspond to the linear, RBF, and polynomial kernels (from top to bottom). When the KPCA is applied, a tensor, *x*_*ijk*_, is unfolded to a matrix, 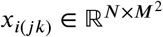.

The performance evaluation is realized as follows. For the KTD, *u*_*𝓁 j*_ *u*_*𝓁 k*_, 1 ≤ *𝓁* ≤ *M*, are divided into two classes, 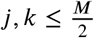,or others, for each *𝓁*. A two way *t* test is performed on these classes. The computed *M P* -values for each *𝓁* are corrected considering the BH criterion (Taguchi, 2020), and the smallest *P* -value is recorded. This process is repeated 100 times, while generating new random variables from the Gaussian distribution. The geometric mean of the *P* -value is computed as a performance measure. For the KPCA, *u*_*𝓁*(*jk*)_, 1 ≤ *𝓁* ≤ *M*^2^ are divided into two classes for each *𝓁*, and the computed *M*^2^ *P* -values for each *𝓁* are corrected using BH criterion. The smallest *P* -value is recorded. This process is repeated 100 times, while generating new random variables from the Gaussian distribution used to generate *x*_*ijk*_. The geometric mean of the *P* -value is computed as a performance measure. Table 1 presents the results of this analysis. It can be noted that the raw *P* -values for the KPCA and KTD are not considerably different; nevertheless, the corrected *P* -values are smaller in the KTD than KPCA for the RBF and linear kernel. In other words, the KTD can effectively identify the singular-value vectors coincident in the two classes with a smaller number of singular-value vectors. Nevertheless, the RBF kernel could not outperform the linear kernel. We implemented other *α* values for the RBF kernel, but the RBF kernel could not outperform the linear kernel through any of the *α* values. This finding suggests that in *large p small n* situations, the introduction of nonlinearity in the kernel is not entirely beneficial, which is likely why the TD-based unsupervised FE could outperform the conventional statistical methods despite its linearity. Furthermore, the introduction of the nonlinearity in the kernels could not improve the performance in *large p small n* situations even when the KTD-based unsupervised FE was applied to real problems, as described in the following sections.

**Table 1.** Geometric mean *P* -values computed through the *t* tests performed on singular-value vectors determined using the KTD and KPCA. Smaller *P* -values are better. *N* = 1000, *N*_1_ = 10, *M* = 6, *µ* = 2, *σ* = 1.

| Kernel<br>$P$-values | RBF ($\alpha = 10^{-6}$) | | RBF ($\alpha = 10^{-3}$) | | linear | |
| --- | --- | --- | --- | --- | --- | --- |
|  | raw | corrected | raw | corrected | raw | corrected |
| KTD | $8.19 \times 10^{-3}$ | $4.31 \times 10^{-2}$ | $1.95 \times 10^{-2}$ | $9.22 \times 10^{-2}$ | $7.18 \times 10^{-3}$ | $3.87 \times 10^{-2}$ |
| KPCA | $9.45 \times 10^{-3}$ | $2.59 \times 10^{-1}$ | $1.45 \times 10^{-2}$ | $3.89 \times 10^{-1}$ | $7.30 \times 10^{-3}$ | $2.01 \times 10^{-1}$ |
| Kernel<br>$P$-values | polynomial ($d = 2$) | | polynomial ($d = 3$) | | | |
|  | raw | corrected | raw | corrected |  |  |
| KTD | $1.36 \times 10^{-1}$ | $4.38 \times 10^{-1}$ | $3.03 \times 10^{-1}$ | $3.61 \times 10^{-1}$ | | |
| KPCA | $7.43 \times 10^{-2}$ | $6.43 \times 10^{-1}$ | $2.28 \times 10^{-1}$ | $3.81 \times 10^{-1}$ | | |

We also examined if the restriction of *u*_*𝓁*(*jk*)_ to 1 ≤ *𝓁* ≤ *M* could improve the performance of the KPCA owing to the smaller number of *P* -values considered in the results. The performance was not improved, indicating that the smallest *P* -values for the KPCA lie in *𝓁 > M*; thus, *u*_*𝓁*(*jk*)_, *M < 𝓁* ≤ *M*^2^ cannot be neglected.

#### SARS-CoV-2 data set

We applied the KTD-based unsupervised FE to real data sets. The first dataset corresponded to the repurposing of drugs for COVID-19, which the TD-based unsupervised FE was successfully applied to the gene expression profiles of SARS-CoV-2 infected cell lines (Blanco-Melo et al., 2020; Taguchi and Turki, 2020). The TD-based unsupervised FE could predict many promising drugs including ivermectin, the clinical trials of which have been recently initiated. Herein, we briefly summarize the process implemented in the previous work (Taguchi and Turki, 2020) to enable a comparative analysis of the results of the KTD-based unsupervised FE with the previous results. The gene expression profiles were formatted as a tensor, *x*_*ijkm*_ ∈ ℝ^*N*×5×2×3^, which indicated whether the gene expression of the *i*th gene of the *j*th cell line was infected (*k* = 1) or not (*k* = 2, control), considering three biological replicates. The HOSVD was applied to *x*_*ijkm*_ to yield

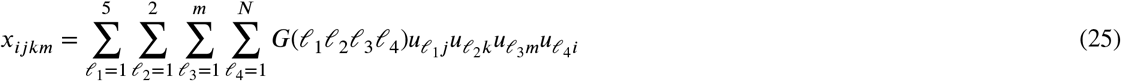

where *G*(*𝓁*_1_*𝓁*_2_*𝓁*_3_*𝓁*_4_) ∈ ℝ ^5×2×3×*N*^ is a core tensor, and 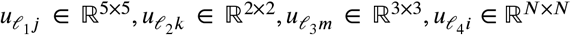 are orthogonal singular value matrices.

The purpose of the analysis was to identify the genes whose expression was distinct between the control and infected cells, independent of the cell lines and replicates. To this end, we selected *𝓁*_1_, *𝓁*_2_, *𝓁*_3_ with constant 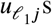 and 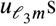 and 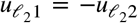. It was noted that *𝓁*_1_ = *𝓁*_3_ = 1 and *𝓁*_2_ = 2 could satisfy these requirements. Next, we identified *𝓁*_4_ = 5 associated with *G* having the largest absolute values given *𝓁*_1_ = 1, *𝓁*_2_ = 2, *𝓁*_3_ = 1. Once *𝓁*_4_ = 5 was selected, the *P* -values were assigned to gene *i*, assuming the null hypothesis that 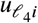 obeys the Gaussian distribution:

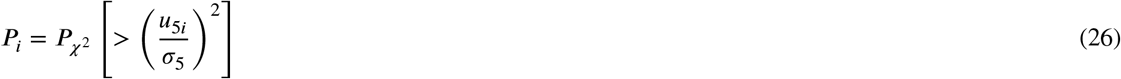

where 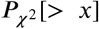 is the cumulative *χ*^2^ distribution in which the argument is larger than *x*, and *σ* _5_ is the standard deviation. The obtained *P* -values were corrected using the BH criterion (Taguchi, 2020). A total of 163 genes for which the adjusted *P* -values were less than 0.01 were selected and used to predict the drugs effective against COVID-19.

The objective of this study was to compare the performance of the KTD-based unsupervised FE applied to *x*_*ijkm*_ with that of an existing study (Taguchi and Turki, 2020). Therefore, we employed the RBF kernel as

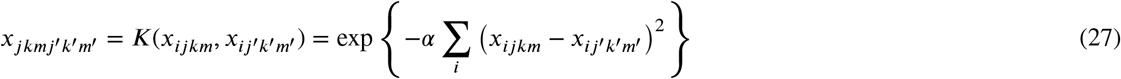

with *α* = 1 × 10^−2^, 1 × 10^−4^, 1 × 10^−6^, to which the HOSVD was applied. We obtained

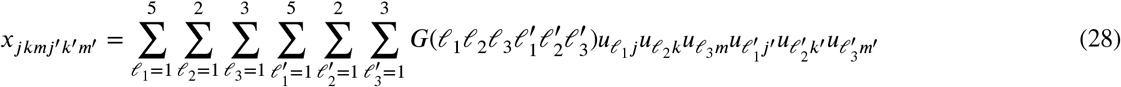

It was observed that *u*_1*j*_ and *u*_1*m*_ were constant, and 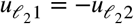 with *𝓁*_2_ = 2. We compared the *u*_1*j*_*u*_2*k*_*u*_1*m*_ computed using TD-based unsupervised FE (Taguchi and Turki, 2020) and KTD with *k*. It was clarified that *u*_1*j*_*u*_2*k*_*u*_1*m*_ computed using the KTD were more notably coincident with *k* (Fig. 3). Thus, the KTD exhibited a slight improvement over the TD-based unsupervised FE (Taguchi and Turki, 2020). Next, we were required to select genes, although the process was challenging as 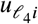 used for gene selection could not be obtained. To apply the KTD for gene selection, we recomputed *u*_1*j*_*u*_2*k*_*u*_1*m*_ excluding *i* sequentially. In addition, the correlation coefficients between *u*_1*j*_*u*_2*k*_*u*_1*m*_ and *k* were obtained. Subsequently, the *i*s were ranked in ascending order of the absolute values of the correlation coefficients. Excluding the genes with expressions coincident with the distinction between *k* = 1 and *k* = 2 was expected to result in less significant (i.e., smaller absolute values of) correlation coefficients. Table 2 presents the confusion matrix of the selected genes between the previous study (Taguchi and Turki, 2020) and KTD. Notably, the genes are highly coincident and can even be regarded as identical considering that there exists 20,000 genes, of which we selected only 163.

**Table 2.** Confusion matrix of the selected genes for the KTD and previous study (Taguchi and Turki, 2020). In the KTD-based unsupervised FE, the ranking was based on the correlation coefficients.

|  |  | KTD-based unsupervised FE |  |  |  |  |  |
| --- | --- | --- | --- | --- | --- | --- | --- |
|  |  | rank |  |  |  |  |  |
| | | $\alpha = 10^{-2}$ | | $\alpha = 10^{-4}$ | | $\alpha = 10^{-6}$ | |
| | | $> 163$ | $\leq 163$ | $> 163$ | $\leq 163$ | $> 163$ | $\leq 163$ |
| Previous study | adjusted $P$-values $\geq 0.01$ | 21516 | 118 | 21509 | 125 | 21540 | 94 |
| | adjusted $P$-values $< 0.01$ | 118 | 45 | 125 | 38 | 94 | 69 |
| Fisher's exact test | $P$-value | $5.05 \times 10^{-59}$ | | $1.53 \times 10^{-46}$ | | $6.61 \times 10^{-108}$ | |
|  | odds ratio | 69 |  | 52 |  | 168 |  |

**Figure 3:**
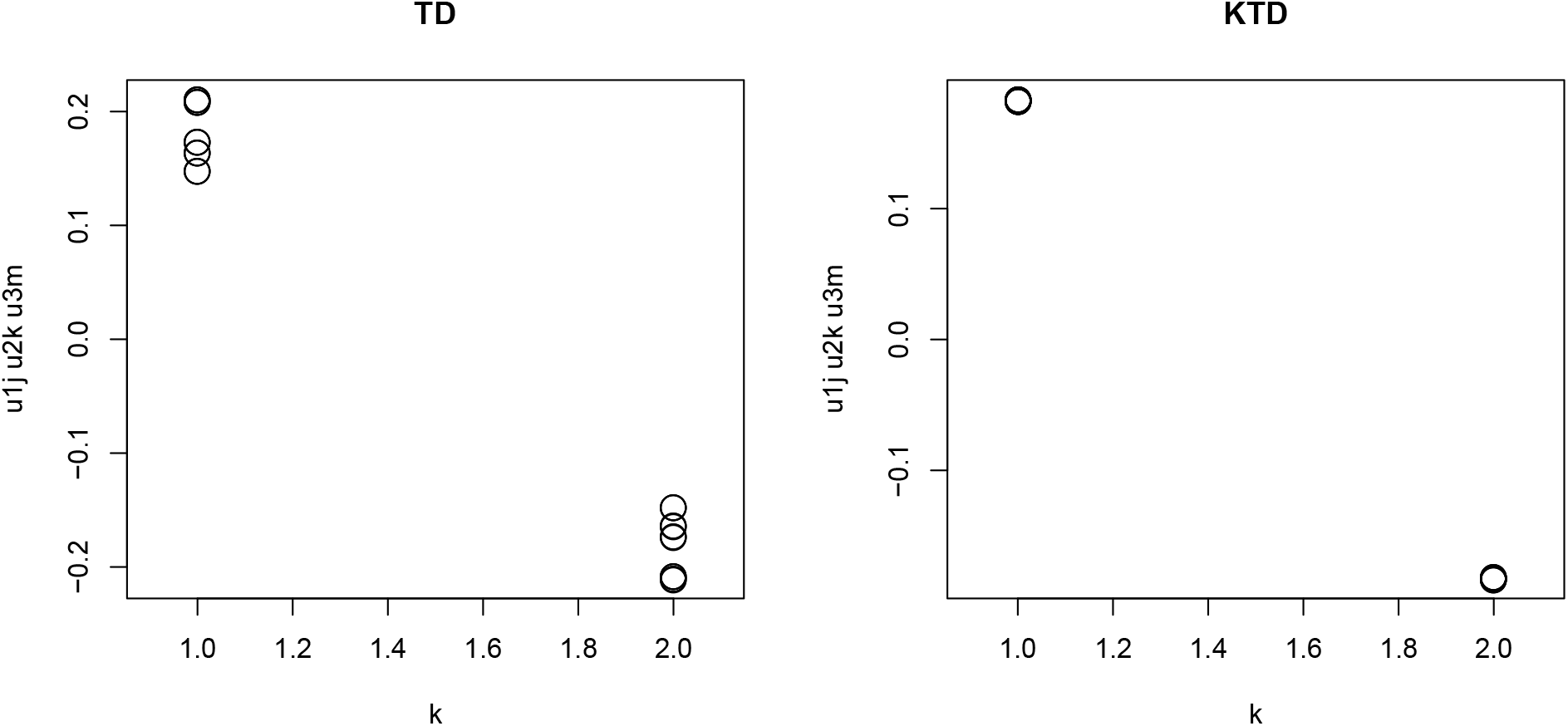
Scatter plot of *u*_1*j*_*u*_2*k*_*u*_1*m*_ and *k*. left:TD (Taguchi and Turki, 2020), right:KTD (*α* = 1 × 10^−6^).

Next, we biologically evaluated the 163 genes selected using the KTD-based unsupervised FE. In particular, we compared the selected 163 genes with gold standard human proteins that are known to interact with the SARS-CoV-2 proteins during infection (Gordon et al., 2020), following the procedure described in the previous study (Taguchi and Turki, 2020) (Table 3).

**Table 3.** Coincidence between 163 genes and human proteins that are known to interact with SARS-CoV-2 proteins during infection. The *P* -values were computed by applying Fisher’s exact tests to the confusion matrix. For more details, please refer to the reference (Taguchi and Turki, 2020).

| SARS-CoV-2<br>proteins | P values |  |  |  | Odds Ratio |  |  |  |
| --- | --- | --- | --- | --- | --- | --- | --- | --- |
| | previous<br>study | $\alpha = 10^{-2}$ | RBF kernel<br>$10^{-4}$ | $10^{-6}$ | previous<br>study | $\alpha = 10^{-2}$ | RBF kernel<br>$10^{-4}$ | $10^{-6}$ |
| SARS-CoV2 E | $6.55 \times 10^{-27}$ | $1.08 \times 10^{-44}$ | $4.90 \times 10^{-22}$ | $6.58 \times 10^{-26}$ | 10.16 | 16.18 | 8.63 | 9.84 |
| SARS-CoV2 M | $1.37 \times 10^{-26}$ | $1.66 \times 10^{-37}$ | $4.14 \times 10^{-22}$ | $4.14 \times 10^{-22}$ | 8.41 | 11.40 | 7.26 | 7.26 |
| SARS-CoV2 N | $4.61 \times 10^{-24}$ | $7.22 \times 10^{-38}$ | $3.51 \times 10^{-15}$ | $1.46 \times 10^{-29}$ | 11.42 | 17.11 | 7.93 | 13.63 |
| SARS-CoV2 nsp1 | $1.05 \times 10^{-20}$ | $3.98 \times 10^{-28}$ | $4.26 \times 10^{-14}$ | $5.51 \times 10^{-15}$ | 10.00 | 12.92 | 7.43 | 7.78 |
| SARS-CoV2 nsp10 | $3.40 \times 10^{-20}$ | $9.04 \times 10^{-26}$ | $4.44 \times 10^{-17}$ | $3.88 \times 10^{-19}$ | 11.51 | 14.13 | 10.06 | 11.02 |
| SARS-CoV2 nsp11 | $1.13 \times 10^{-29}$ | $2.62 \times 10^{-38}$ | $4.00 \times 10^{-20}$ | $1.2 \times 10^{-27}$ | 10.66 | 13.43 | 7.74 | 10.03 |
| SARS-CoV2 nsp12 | $4.87 \times 10^{-20}$ | $1.64 \times 10^{-29}$ | $1.85 \times 10^{-14}$ | $2.41 \times 10^{-15}$ | 9.48 | 13.09 | 7.38 | 7.72 |
| SARS-CoV2 nsp13 | $6.04 \times 10^{-33}$ | $3.08 \times 10^{-37}$ | $2.19 \times 10^{-20}$ | $7.41 \times 10^{-28}$ | 11.17 | 12.49 | 7.51 | 9.65 |
| SARS-CoV2 nsp14 | $1.75 \times 10^{-22}$ | $1.08 \times 10^{-31}$ | $2.72 \times 10^{-16}$ | $2.78 \times 10^{-18}$ | 12.05 | 16.26 | 9.30 | 10.18 |
| SARS-CoV2 nsp15 | $1.85 \times 10^{-20}$ | $4.26 \times 10^{-29}$ | $1.71 \times 10^{-17}$ | $1.08 \times 10^{-14}$ | 10.23 | 13.75 | 9.03 | 7.90 |
| SARS-CoV2 nsp2 | $4.81 \times 10^{-33}$ | $5.16 \times 10^{-42}$ | $3.31 \times 10^{-20}$ | $6.34 \times 10^{-31}$ | 11.79 | 14.76 | 7.83 | 11.11 |
| SARS-CoV2 nsp4 | $5.79 \times 10^{-29}$ | $5.25 \times 10^{-42}$ | $4.09 \times 10^{-16}$ | $5.53 \times 10^{-26}$ | 10.26 | 14.41 | 6.47 | 9.36 |
| SARS-CoV2 nsp5 | $3.78 \times 10^{-25}$ | $4.18 \times 10^{-38}$ | $6.97 \times 10^{-18}$ | $4.63 \times 10^{-24}$ | 12.36 | 17.97 | 9.36 | 11.91 |
| SARS-CoV2 nsp5+ | $3.75 \times 10^{-17}$ | $1.42 \times 10^{-21}$ | $3.74 \times 10^{-17}$ | $4.65 \times 10^{-14}$ | 11.39 | 13.78 | 11.39 | 9.72 |
| SARS-CoV2 nsp6 | $9.47 \times 10^{-26}$ | $7.15 \times 10^{-35}$ | $3.91 \times 10^{-21}$ | $4.97 \times 10^{-22}$ | 9.00 | 11.71 | 7.68 | 7.94 |
| SARS-CoV2 nsp7 | $1.93 \times 10^{-29}$ | $1.09 \times 10^{-33}$ | $2.70 \times 10^{-16}$ | $1.61 \times 10^{-22}$ | 10.81 | 12.18 | 6.76 | 8.65 |
| SARS-CoV2 nsp8 | $1.11 \times 10^{-29}$ | $1.68 \times 10^{-41}$ | $1.57 \times 10^{-18}$ | $1.11 \times 10^{-28}$ | 10.14 | 13.80 | 6.95 | 9.84 |
| SARS-CoV2 nsp9 | $5.36 \times 10^{-29}$ | $1.46 \times 10^{-40}$ | $8.17 \times 10^{-19}$ | $6.71 \times 10^{-28}$ | 12.24 | 16.67 | 8.59 | 11.85 |
| SARS-CoV2 orf10 | $5.29 \times 10^{-34}$ | $2.44 \times 10^{-44}$ | $5.57 \times 10^{-20}$ | $1.16 \times 10^{-26}$ | 12.37 | 15.95 | 7.94 | 10.01 |
| SARS-CoV2 orf3a | $2.06 \times 10^{-28}$ | $1.31 \times 10^{-43}$ | $3.17 \times 10^{-18}$ | $1.62 \times 10^{-24}$ | 9.95 | 14.76 | 6.99 | 8.80 |
| SARS-CoV2 orf3b | $1.89 \times 10^{-29}$ | $7.58 \times 10^{-41}$ | $1.35 \times 10^{-17}$ | $2.50 \times 10^{-27}$ | 11.80 | 15.94 | 7.78 | 11.06 |
| SARS-CoV2 orf6 | $8.81 \times 10^{-26}$ | $5.67 \times 10^{-39}$ | $5.22 \times 10^{-13}$ | $8.03 \times 10^{-22}$ | 10.37 | 14.99 | 6.14 | 9.05 |
| SARS-CoV2 orf7a | $1.69 \times 10^{-28}$ | $9.79 \times 10^{-35}$ | $8.26 \times 10^{-22}$ | $1.18 \times 10^{-23}$ | 10.00 | 11.90 | 8.04 | 8.57 |
| SARS-CoV2 orf8 | $5.94 \times 10^{-28}$ | $2.48 \times 10^{-39}$ | $4.32 \times 10^{-18}$ | $1.27 \times 10^{-20}$ | 9.25 | 12.57 | 6.57 | 7.25 |
| SARS-CoV2 orf9b | $6.54 \times 10^{-30}$ | $7.07 \times 10^{-44}$ | $5.12 \times 10^{-17}$ | $9.11 \times 10^{-28}$ | 12.12 | 17.34 | 7.68 | 11.36 |
| SARS-CoV2 orf9c | $1.11 \times 10^{-28}$ | $1.03 \times 10^{-42}$ | $1.15 \times 10^{-21}$ | $5.76 \times 10^{-26}$ | 8.35 | 12.07 | 6.66 | 7.69 |
| SARS-CoV2 Spike | $8.22 \times 10^{-26}$ | $3.48 \times 10^{-41}$ | $8.46 \times 10^{-15}$ | $6.07 \times 10^{-22}$ | 10.08 | 15.34 | 6.55 | 8.81 |

+ with mutation, C145A

The 163 genes selected by the KTD-based unsupervised FE are highly coincident with the gold standard SARS-CoV-2 interacting human proteins, the results in Table 3 even outperformed the results for the gold standard SARS-CoV-2 interacting human proteins compared to the results presented in Table S33 (Taguchi and Turki, 2020) corresponding to the application of the TD-based unsupervised FE (Taguchi and Turki, 2020). Notably, all 27 protein groups when *α* = 10^−2^ exhibit larger odds ratio as well as smaller *P* -values than those in the previous study (corresponding to the TD-based unsupervised FE (Taguchi and Turki, 2020)) in Table 3.

Tables 3 indicates that the KTD based unsupervised FE outperformed the TD-based unsupervised FE when it was applied to a real data set in contrast to when the KTD-based unsupervised FE was applied to synthetic data simulating a *large p small n* problem (eq.(21)). This is possibly because synthetic data set could not fully emulate complicated structures in real data sets; KTD-based unsupervised FE should be tested with more real data sets to test whether it can outperform TD-based unsupervised FE.

#### Kidney cancer data sets

In a previous work, we attempted to realize the integrated analysis of multiomics data by applying the TD-based unsupervised FE (Ng and Taguchi, 2020). In the existing study (Ng and Taguchi, 2020), the objective was to select genes that act as prognostic biomarker for kidney cancer. To this end, the TD-based unsupervised FE was employed to identify the differentially expressed genes (DEGs) between normal kidneys and tumors in two independent data sets. The TD-based unsupervised FE successfully identified 72 mRNAs and 11 miRNAs for the first data set retrieved from the TCGA, and 209 mRNAs and 3 miRNAs for the second data set retrieved from the GEO. To extend this analysis, in the present study, the mRNA expression matrix *x*_*ij*_ ∈ ℝ^*N*×*M*^, which represents the *i*th mRNA expression in the *j*th sample and the miRNA expression matrix *x*_*kj*_ ∈ ℝ^*K*×*M*^, which represents the *k*th miRNA expression in the *j*th sample were integrated into the tensor *x*_*ijk*_ ∈ ℝ^*N*×*M*×*K*^ as

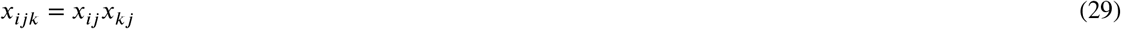

and converted to a matrix *x*_*ik*_ ∈ ℝ^*K*×*M*^

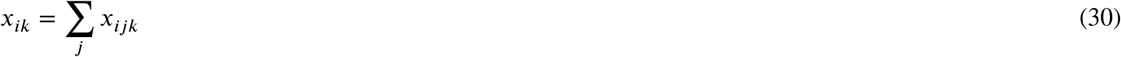

to which the SVD was applied to yield

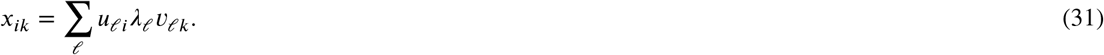

The missing singular-value vectors attributed to the *j*th mRNA and miRNA samples were recovered as follows:

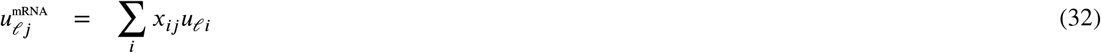

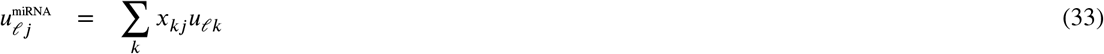

The *P* -values attributed to mRNAs and miRNAs were derived as

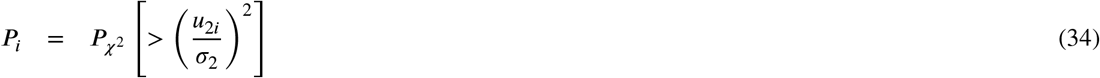

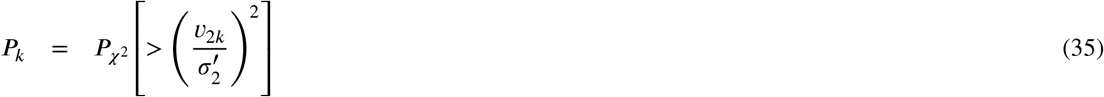

as 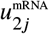 and 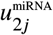 were noted to be coincident with the distinction between the tumors and normal kidneys. The mRN^2*j*^As and m^2^i^*j*^RNAs with the adjusted *P* -values of less than 0.01 were selected.

Notably, the kernel trick cannot be applied to this formulation as the summation is obtained over the sample index *j* and not the feature index *i, k*. Thus, instead, we formulated a kernel-trick-friendly integration of *x*_*ij*_ and *x*_*kj*_ as

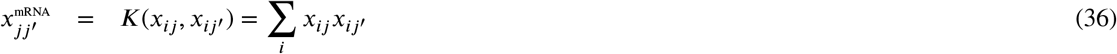

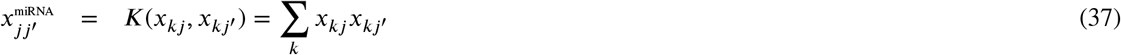

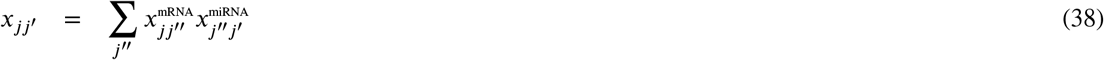

to which the SVD can be applied to yield

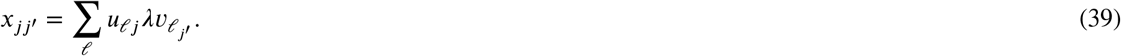

As this expression includes the inner product, for which the summation is obtained over the feature index *i* (Eq. (36)) and *k* (Eq. (37)), the expressions can be easily extended to the RBF kernel as

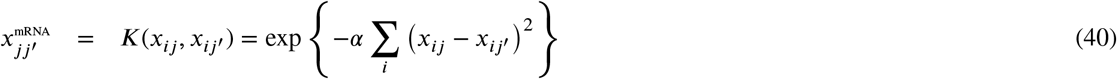

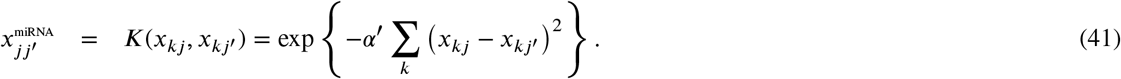

*x*_*jj*_′ was computed using Eq. (38), and the SVD was applied to *x*_*jj*_′ to obtain an expression similar to Eq. (39).

A possible biological validation of the performances of the KTD-based unsupervised FE is by evaluating *u*_*𝓁j*_ and *v*_*𝓁j*_, corresponding to the mRNA and miRNA samples, respectively. Specifically, we consider whether

- *u*_*𝓁j*_s are distinct between tumorous and normal kidneys,
- *v*_*𝓁j*_s are distinct between tumorous and normal kidneys,
- *u*_*𝓁j*_s and *v*_*𝓁j*_s are coincident.

To examine the first point, we compute *u*_*𝓁j*_ and *v*_*𝓁j*_ using linear (eqs. (36) and (37)) and RBF (eqs. (40) and (41)) kernels. As in the previous study (Ng and Taguchi, 2020), the second singular-value vectors, *u*_2*j*_ and *v*_2*j*_, are noted to be the most coincident with the aforementioned three requirements. Table 4 presents the comparisons of the *P* –values used to evaluate the aforementioned three conditions for the linear kernel, RBF kernel, and the approach employed in the previous study. The *P* -values to evaluate the distinction between tumorous and normal kidneys were computed through the *t* test, whereas those evaluating the coincidence between *u*_2*j*_ and *v*_2*j*_ were computed by considering the correlation coefficients. Although the three types of approaches exhibited a reasonable performance, the RBF kernel that considered the nonlinearity could not always outperform the two other linear methods. The superiority of the RBF kernel was only noted occasionally depending on *α* and *α* ^′^. Thus, the nonlinear method could not always outperform the linear methods in the *large p small n* situation.

**Table 4.** Performances of the linear and RBF kernel TD-based unsupervised FE and that achieved in the previous study (Ng and Taguchi, 2020) by using Eq. (30). The *P* -values marked by an asterisk correspond to the most significant values.

|  | tumor vs. normal kidney |  | correlation coefficient |
| --- | --- | --- | --- |
| | $u_{2j}$ (mRNA) | $v_{2j}$ (miRNA) | $u_{2j}$ vs. $v_{2j}$ |
| 1st data set (TCGA) |  |  |  |
| linear kernel | $1.27 \times 10^{-38}$ | $1.76 \times 10^{-62}$ | 0.912 ($1.28 \times 10^{-126}$) |
| RBF kernel ($\alpha = 10^{-10}$ and $\alpha' = 10^0$) | $1.99 \times 10^{-14}$ | $1.99 \times 10^{-14}$ | 1.00 (0.00*) |
| RBF kernel ($\alpha = 10^{-12}$ and $\alpha' = 10^{-2}$) | $1.06 \times 10^{-44}$* | $4.24 \times 10^{-38}$ | 0.974 ($5.93 \times 10^{-212}$) |
| RBF kernel ($\alpha = 10^{-14}$ and $\alpha' = 10^{-4}$) | $4.70 \times 10^{-43}$ | $2.89 \times 10^{-66}$ | 0.914 ($2.00 \times 10^{-128}$) |
| previous study | $7.10 \times 10^{-39}$ | $2.13 \times 10^{-71}$* | 0.905 ($1.63 \times 10^{-121}$) |
| 2nd data set (GEO) |  |  |  |
| linear kernel | $5.36 \times 10^{-23}$* | $2.76 \times 10^{-18}$ | 0.937 ($3.91 \times 10^{-16}$) |
| RBF kernel ($\alpha = 10^{-4}$ and $\alpha' = 10^0$) | $5.61 \times 10^{-7}$ | $5.61 \times 10^{-7}$ | 1.00 (0.00*) |
| RBF kernel ($\alpha = 10^{-6}$ and $\alpha' = 10^{-2}$) | $5.42 \times 10^{-15}$ | $1.17 \times 10^{-13}$ | 0.966 ($2.06 \times 10^{-20}$) |
| RBF kernel ($\alpha = 10^{-8}$ and $\alpha' = 10^{-4}$) | $6.93 \times 10^{-18}$ | $2.93 \times 10^{-15}$ | 0.921 ($1.16 \times 10^{-14}$) |
| previous study | $6.74 \times 10^{-22}$ | $2.54 \times 10^{-18}$* | 0.931 ($1.58 \times 10^{-15}$) |

Finally, we compared the genes selected using the three methods. For the RBF, we followed the procedure pertaining to the RBF kernel TD-based unsupervised FE applied to the SARS-CoV-2 infection cell lines *x*_*jj*_′. was computed excluding one of the miRNAs or mRNAs, and the SVD was applied to the obtained *x*_*jj*_′ to obtain *u*_2*j*_ The *t* test was applied to *u*_2*j*_ to compute the *P* -values using which the distinction of *u*_2*j*_ between tumorous and normal kidneys could be evaluated. When the first data set was considered, 72 top ranked mRNAs and 11 top ranked miRNAs having large (thus, less significant) *P* -values were selected because the exclusion of the mRNAs or miRNAs distinct between the tumorous and normal kidneys was expected to decrease the significance of the distinct *u*_2*j*_ between the two entities. Similarly, when the second data set was considered, 209 top ranked mRNAs and 3 top ranked miRNAs were selected.

For the linear kernel, the *P* -values attributed to the miRNAs and miRNAs were computed using Eqs. (34) and (35) with

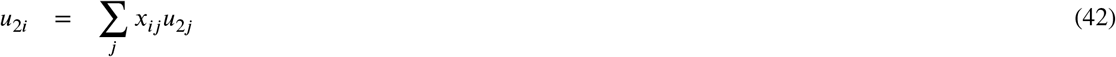

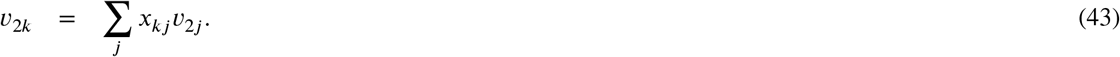

Subsequently, the mRNAs and miRNAs with adjusted *P* -values of less than 0.01 were selected. Table 5 presents a comparison of the confusion matrices of the selected genes between the present study and previous study (Ng and Taguchi, 2020). The higher coincidence of the RBF or linear kernel with that obtained in the previous study depended on *α* and *α*^′^. The RBF kernels were more coincident with the findings of the previous study for smaller *α* and *α*^′^, although liner kernel was always more coincident with the previous study where TD-based unsupervised FE was employed.

**Table 5.**
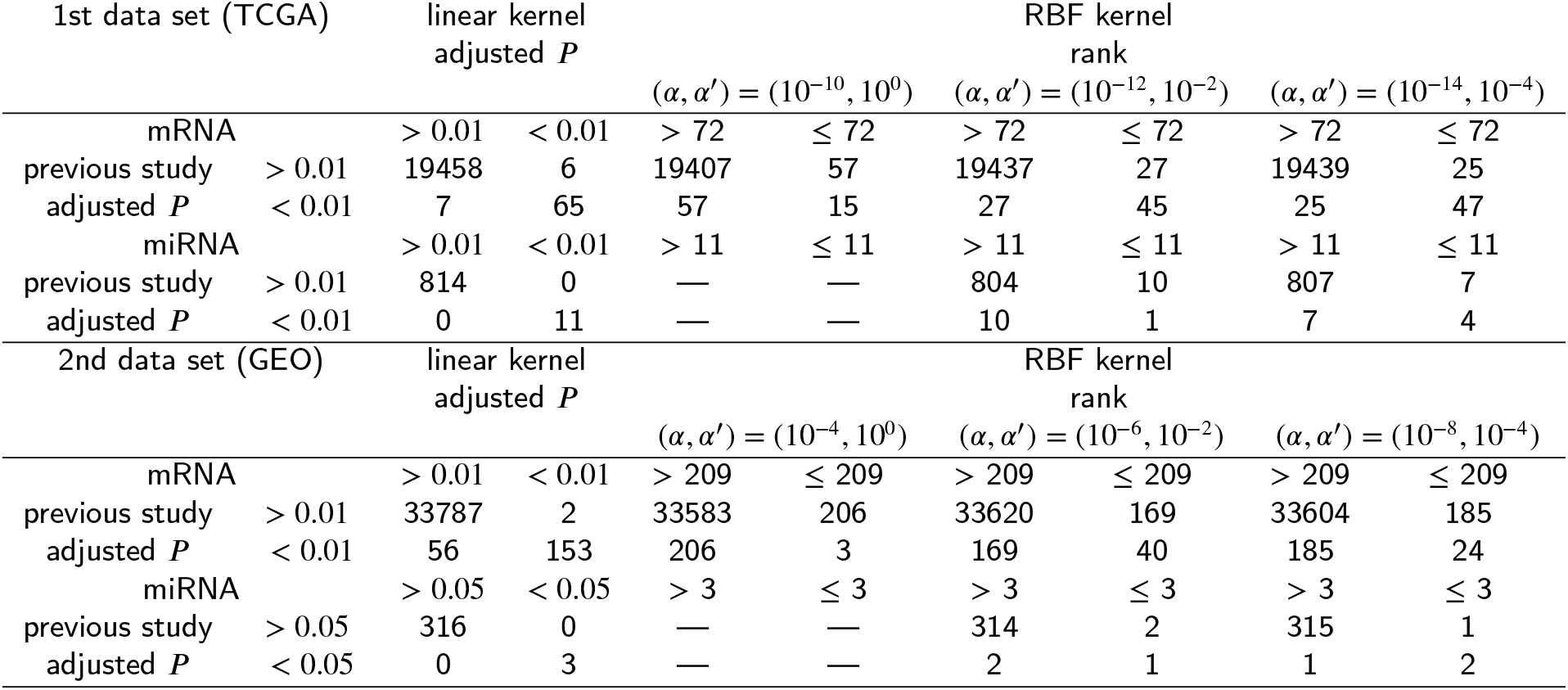
Confusion matrix of the selected genes between the KTD (linear and RBF kernel) and TD-based unsupervised FE (previous study (Ng and Taguchi, 2020) using Eq. (30)). The ranking in the KTD-based unsupervised FE was *t* test based.

Although no gold standard data sets exist to evaluate the selected genes, one possible evaluation pertains to the coincidence of the selected genes between the first (TCGA) and second data sets (GEO). As these two data sets are independent, it is unlikely that commonly selected genes are present, and the selected genes are at most 1% of all genes. In the previous study (Ng and Taguchi, 2020) that employed Eq. (30), 11 genes were commonly selected between TCGA and GEO, whereas as small as 7 genes were commonly selected between the TCGA and GEO for the linear and RBF kernels (*α* = 10^−12^ and *α*^′^ = 10^−2^ that achieved the maximum overlap among the tested RBF kernels). Although these overlaps are still significant (using Fisher’s exact test, *P* = 3.43 × 10^−6^, odds ratio: 12.54 for RBF kernel and *P* = 7.93 × 10^−7^, odds ratio: 15.82 for linear kernel), the values are inferior to those for the previous study (Ng and Taguchi, 2020) that involved 11 commonly selected genes (*P* = 8.97 × 10^−11^, odds ratio: 19.7). Thus, the KTD-based unsupervised FEs are at most comparable with the TD-based unsupervised FE in this *large p small n* problem.

## Discussion

In the aforementioned analyses, we successfully formalized the KTD-based unsupervised FE to be applied to select DEGs. Although the implementation was effective, the KTD-based unsupervised FE could not always outperform the TD based unsupervised FE that address only the linearities. This finding may appear to be unexpected, as nonlinear methods generally outperform linear methods pertaining to nonlinear application by definition. Nevertheless, this expectation does not always hold, especially when the linear methods can achieve optimal solutions similar to those attained by nonlinear methods.

As shown in Fig. 2(D), the KTD can precisely recognize the nonlinearity in Swiss Rolls, at least partially. However, when the KTD-based unsupervised FE is applied to *large p small n* problems, even the use of nonlinear kernels could not always outperform the TD-based unsupervised FE or linear kernel based KTD-based unsupervised FE. The reason for this aspect can be explained as follows. To represent complicated structures that only nonlinear methods can recognize, many points are required. For example, to represent *n*th order polynomials, we need at least *n* + 1 points. As the higher order polynomials can represent more complicated structures, more points are necessary to realize this representation. In *large p small n* problems, the number of samples is insufficient to represent complicated structures. As shown in Fig. 2, there exist 1,000 points in 3-dimensional space. In contrast, in the considered *large p small n* problems, the number of samples ranges from a few tens to hundreds, whereas the number of corresponding features varies from 10^2^ to 10^4^. Thus, clearly, the points cannot effectively represent complicated structures. The TD-based unsupervised FE, despite its linearity, has outperformed conventional statistical test-based gene selection methods. It is rather unexpected that such a simple linear method can outperform more advanced tools. Specifically, the TD-based unsupervised FE has already achieved the performances that KTD-based unsupervised FE employing nonlinear kernels is expected to achieve in *large p small n* problems.

In addition, the KTD-based unsupervised FE cannot exploit certain advantages that the TD-based unsupervised FE exhibits. For example, as the KTD cannot yield singular-value vectors attributed to the features (genes), we cannot apply the empirical *P* -value computation assuming that the singular-value vectors obey the Gaussian distribution. Instead, we must repeatedly compute the singular-value vectors excluding the features sequentially, which is a time-consuming process. In addition, the computation of inner products exponentially doubles the amount of memory required. As the number of samples is small, this aspect may not be an immediate problem. Nevertheless, this process for the first data set (TCGA) required half a day of CPU time, whereas using 12 CPU units as the set included hundreds of samples. In contrast, the TD-based unsupervised FE can complete the corresponding computation in a few minutes. Furthermore, the *P* -values computed by the KTD-based unsupervised FE cannot be directly used for gene selection. When used to rank the genes, the process is effective (Tables 2 and 5). However, the process is ineffective when used to screen features directly because the *P* -values are sufficiently small even after excluding one variable feature, e.g., those distinct between two classes (e.g., control and cancers). In this scenario, we cannot identify the number of features to be selected by only considering the *P* -values. Nevertheless, this decision can be made using TD-based unsupervised FE.

Despite these disadvantages of KTD-based unsupervised FE, there may exist scenarios in which the results can be improved, whereas this aspect does not hold true for TD-based unsupervised FE that lacks tunable parameters. At present, although we were unable to identify the cases in which the KTD unsupervised FE employing nonlinear kernels could outperform the TD-based unsupervised FE in *large p small n* problems, it is expected that the proposed approach can outperform the existing approach in scenarios involving *large p small n* problems in which the TD-based unsupervised FE cannot achieve a high performance.

The reason we reviewed neither recent approaches dealing with *large p small n* problems nor tensor learning approaches is primarily because our approach is very distinct from other approaches. First, the recent approaches that deal with *large p small n* problems are not applicable to the extreme cases that we dealt with in this study. In this study, the numbers of features (i.e., genes) are 10^4^, whereas the numbers of samples are from 10 to 10^2^. Thus, the ratio of the former to the latter is as large as 10^2^, whereas conventional approaches regarded to be applicable to *large p small n* problems usually do not assume such extreme cases. For example, although Loh (2011) reviewed various tools that deal with *large p small n* problems, he applied the methods to at most *p* = 500 whereas *n* = 100; the number of features is only 5 times larger than that of samples, which is much less than 10^2^. Chakraborty et al. (2012) applied Bayesian nonlinear regression for *large p small n* problems that have only 700 features for which as many as 62 samples were measured; thus, the ratio of the number of features to the number of samples is about 10, which is also much less than 10^2^. Secchi et al. (2013) dealt with extreme case where *n* = 20 and *p* = 1024, whereas they did not apply to feature selection because their simulated data was not composed of the mixture of features with and without distinction between two classes but was composed of features distinct between two classes or those without distinction. Although there are only a few examples and it is impossible to prove that there are no methods applicable to the present data sets, we need the methods satisfying the following conditions:

- applicable to *large p small n* problems where 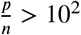
- need to extract generated futures that can discriminate between multiple classes
- selecting a small number of features biologically significant (e.g. enrichment analysis, see Table 3) with stability (e.g., coincidence between two independent data sets, see Table 5)

which are not likely fulfilled with the methods that are not specifically designed to be fitted to genomic science.

This difference between our approach and general machine learning solutions that aim to solve the *large p small n* problem prevents us from applying other methods to our problem. For example, fast adaptive K-means (Wang et al., 2019a) (FAKM) aims to select features coincident with clustering using K-means algorithm. When FAKM was applied to synthetic data (Eq. (21)), in spite of FAKM almost always correctly selecting features *i* ≤*N*_1_, K-means practically fails to generate two clusters coincident with the distinction between 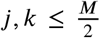 and others. This is because simply selecting *i ≤ N*_1_ features cannot allow K-means to generate correct clusters beca^2^use of the too-noisy data set. Because FAKM simply tries to select features relatively coincident with clustering, it cannot guarantee the generation of the correct clustering. On the other hand, although Wang et al. (2019b) proposed methods that can generate low rank latent variables that can discriminate clusters correctly, it does not have ability to select features. To our knowledge, no other general machine learning methods applicable to the present problem exist.

As for tensor learning, the term “tensor learning” often simply means deriving low rank representations of given tensors, typically using tensor decomposition (Guo et al., 2012; Tao et al., 2007; Liu et al., 2016). Thus, the main purpose of the so-called tensor learning is to approximate the whole tensor with low rank representation. In contrast to these purposes of tensor learning, we need to find latent variables coincident with the properties of interest (e.g., distinction between infectious and control cell lines, Fig. 3), regardless of how well it can approximate a whole tensor. Thus, our purpose differs from the general purpose of tensor learning in which how well it can approximate the whole tensor is the question. From this point view, tensor learning is not deeply related to our above purpose.

As for the comparisons with the methods designed to be fitted to genomic science, superiority of previously proposed TD-based unsupervised FE over the conventional methods was repeatedly demonstrated. For example, when we previously investigated SARS-CoV-2 data set (Taguchi and Turki, 2020) analyzed in this study, three conventional methods frequently used to identify differentially expressed genes, *t* test, SAM (Tusher et al., 2001) and limma (Ritchie et al., 2015) were shown to be inferior to TD-based unsupervised FE. When we previous analyzed kidney cancer data sets (Ng and Taguchi, 2020) analyzed in this study, these three methods were shown to be inferior to TD-based unsupervised FE. Other than these two studies, superiority of TD-based unsupervised FE over conventional methods was repeatedly demonstrated. For example, when TD-based unsupervised FE was applied to gene expression profiles measured in multiple mice tissues under the multiple experimental conditions (Taguchi, 2017), categorical regression (also known as ANOVA), SAM and limma were shown to be inferior to TD-based unsupervised FE. Based on these previous studies, the superiority of TD-based unsupervised FE over conventional methods was established. Therefore, we concentrated on the comparisons between newly proposed kernel TD-based unsupervised FE and original TD-based unsupervised FE in this study; extensive comparisons with other methods than TD-based unsupervised FE was not critical.

Finally, we added more comments about cpu time used for KTD-based unsupervised FE. When liner kernel, Eq. (22) was employed, cpu time was as short as TD-based unsupervised FE because we made use of equations like eqs. (42) and (43) for selecting genes. Nevertheless, when we employ RBF kernel, we need to recompute TD every time we exclude one feature. When we performed one run of RBF kernel based KTD based unsupervised FE to SARS-CoV-2 infection data set, it took 30 minutes on cpu E5-2643 3.40GHz with 256GB memory. One run of the same computation to TCGA kidney data took 300 h on the same machine. CPU time is mainly dependent upon the square of sample size. For SARS-CoV-2, there are only 30 samples, whereas for TCGA kidney data, there are 324 samples. The ratio of samples size is 324/30 ∼10, whose square is 10^2^. The ratio of cpu time required is 300/0.5 ∼ 6 × 10^2^. These 2 match with each other excluding numerical factor less than 10. Thus, it is unrealistic to apply KTD-based unsupervised FE to the samples larger than 10^3^.

## Methods

### Data set

The data sets used in this study are all public domain. The information regarding the retrieval of these data sets has been presented in previous studies (Taguchi and Turki, 2020; Ng and Taguchi, 2020), in which the TD-based unsupervised FE approach was applied to these data sets.

## Acknowledgements

This work was supported by KAKENHI [grant numbers 19H05270, 20H04848, and 20K12067] to YT and Deanship of Scientific Research (DSR) at King Abdulaziz University, Jeddah [grant number KEP-8-611-38] to TT.

## Author contributions statement

YHT: Conceptualization, Formal analysis, Methodology, Software, Supervision, Writing – original draft, Writing – review & editing. TT: Data curation, Writing – original draft, Writing – review & editing

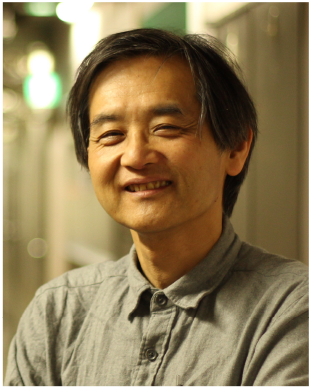

Y-H. Taguchi received a B.S. degree in physics from the Tokyo Institute of Technology and a Ph.D. degree in physics from the Tokyo Institute of Technology. He is currently a full professor with the Department of Physics, Chuo University, Japan. His works have been published in leading journals such as Physical Review Letters, Bioinformatics, and Scientific Reports. His research interests include bioinformatics, machine-learning, and nonlinear physics. He is also an editorial board member of Frontiers in Genetics:RNA, PloS ONE, BMC Medical Genomics, Medicine (Lippincott Williams & Wilkins journal), BMC Research Notes, noncoding RNA (MDPI), and IPSJ Transaction on Bioinformatics.

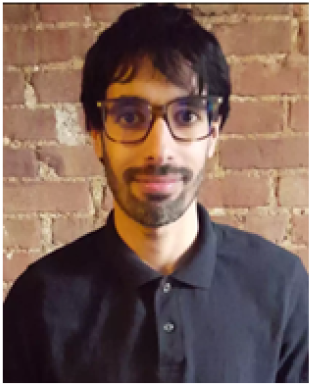

Turki Turki received a B.S. degree in computer science from King AbdulAziz University, an M.S. degree in computer science from NYU.POLY, and a Ph.D. degree in computer science from the New Jersey Institute of Technology. He is currently an assistant professor with the Department of Computer Science, King Abdulaziz University, Saudi Arabia. His research interests include Artificial Intelligence (Tensor Learning, Machine Learning, Deep Learning) and Bioinformatics. His research studies have been published in journals such as Expert Systems with Applications, Frontiers in Genetics, Current Pharmaceutical Design, Computers in Biology and Medicine, and Genes. Dr. Turki has served on the program committees of several international conferences and is currently a review editor for Frontiers in Artificial Intelligence and Frontiers in Big Data. In addition, he is an editorial board member of Computers in Biology and Medicine and Sustainable Computing: Informatics and Systems.

